# Individual- and population-level drivers of consistent foraging success across environments

**DOI:** 10.1101/260604

**Authors:** Lysanne Snijders, Ralf H. J. M. Kurvers, Stefan Krause, Indar W. Ramnarine, Jens Krause

**Affiliations:** Department of Biology and Ecology of Fishes, Leibniz-Institute of Freshwater Ecology and Inland Fisheries, Müggelseedamm 310, 12587 Berlin, Germany; Center for Adaptive Rationality, Max Planck Institute for Human Development, Lentzeallee 94, 14195 Berlin, Germany; Department of Electrical Engineering and Computer Science, Lübeck University of Applied Sciences, Mönkhofer Weg 239, 23562 Lübeck, Germany; Department of Life Sciences, University of the West Indies, St Augustine, Trinidad and Tobago; Faculty of Life Sciences, Humboldt-Universitӓt zu Berlin, Invalidenstrasse 42, 10115 Berlin, Germany

## Abstract

Individual foraging is under strong natural selection. Yet, whether individuals differ consistently in their foraging success across environments, and which individual and population-level traits might drive such differences, is largely unknown. We addressed this question in a field experiment, conducting over 1,100 foraging trials with nine subpopulations of guppies, *Poecilia reticulata*, translocating them across environments in the wild. A-priori, we determined the individual social phenotypes. We show that individuals consistently differed in reaching food, but not control, patches across environments. Social individuals reached more food patches than less social ones and males reached more food patches than females. Overall, individuals were, however, more likely to join females at patches than males, which explains why individuals in subpopulations with relatively more females reached, on average, more food patches. Our results provide rare evidence for individual differences in foraging success across environments, driven by individual and population level (sex ratio) traits.

---

Animals strongly depend on successful foraging (e.g. the localization and acquisition of food resources) for their survival and reproduction and this success is, in turn, shaped by the environment. Often individuals have to locate resources that vary unpredictably in their distribution through space (patchy) and time (sporadic)^1,2^. Next to varying resource distributions, individuals regularly see the environment itself (or their experience of it) change over time, for example due to a fluctuating lunar cycles^3^, seasonality^4^ or dispersal and migration initiated by the animals themselves^5,6^. Animals living in unpredictable and changing environments can thus be expected to possess traits that enable them to maintain consistent foraging success under such uncertain conditions.

Evidence that individual foraging behaviour in the wild can be consistent over time is accumulating^7,8^, but evidence for consistency of foraging behaviour across environments remains rare and evidence for consistency of foraging success across environments is, to our knowledge, absent. Yet, it is crucial to study individuals in different environments if we are to disentangle the causal factors of foraging success. Moreover, when we know if individuals consistently differ in finding unpredictable resources in fluctuating environments and identify which individual traits underlie this ability, we can better understand how natural selection might act upon such traits.

Here we combined over 1,100 ecologically realistic foraging trials with dynamic social (Markov Chain) modelling^9^ in nine subpopulations of guppies, *Poecilia reticulata*, to address whether foraging success indeed varies consistently between individuals across different environments in the wild and whether individual traits, such as sex and social tendency, explain such variation. To measure social tendency, we quantified the social dynamics of each subpopulation via focal observations prior to the foraging trials. During the foraging trials, we presented food items (‘patches’ from now on) at various locations in a novel environment (pool) and determined the identity, order and feeding behaviour of each fish that arrived within one minute of the first arriving fish. We examined consistency of foraging success, in terms of the localization of novel food patches across environments, by translocating entire subpopulations between different natural pools. Furthermore, we accounted for success driven merely by individual differences in movement behaviour, by additionally testing individual responses towards unpredictably distributed control patches (i.e. items without food).

Guppies are an ideal study system to address these fundamental questions regarding foraging ecology. First, they inhabit rainforest rivers that change dynamically throughout the year, separating subpopulations in temporarily isolated pools during the dry season. In here, these fish forage on items that are patchily and sporadically distributed through time and space. Second, dynamic social network analyses have revealed that guppies are consistent in their social tendencies (i.e. time spent social) across different environments in the wild^10,11^. Overall, social tendency does not differ between the sexes, but males distribute their social contact moments more evenly over social partners than females^11^. Finally, there is strong evidence for socially mediated foraging in, especially female, guppies^12–15^. We therefore expected females and more social individuals to be consistently better foragers across environments.

## Results

In total, 92% of all the presented patches (89% control, 94% food) were ‘detected’, i.e. reached by at least one fish of a subpopulation (‘batch’ from here on). A batch was overall significantly, yet only slightly, more likely to detect a food patch than a control patch (*χ^2^* = 10.99, *P* < 0.001, *Relative Risk* = 1.05 (1.03-1.06), *N* = 1,135). Likelihood of food over control patch detection did not change over time (Treatment*Time: *χ^2^* = 0.10, *P* = 0.75, *N* = 1,135). Control patches thus mimicked food patches very closely, albeit not perfectly (see also Supplementary Fig. 1). For the remainder of the Results section we only consider detected patches (i.e. the 92% of the patches with the potential to provide social information).

### Consistent individual foraging success across environments

Individuals that reached a higher proportion of food patches gained a higher number of foraging bites (*χ^2^* = 100.10, *P* < 0.001, *N* = 114; Supplementary Fig. 2), demonstrating that patch visitation, i.e. localization of resources, is an important component of foraging success. Individuals consistently differed in the proportion of food patches they reached across different pools (Repeatability (*R*) = 0.34, *SE* = 0.13, *CI* = 0.09 – 0.59, *P* = 0.02, *N* = 114; Fig. 1). That is, individuals that reached more food patches in their initial pool also reached more patches after being translocated to another pool. Individuals, however, did not differ in the proportion of control patches they reached, even though these patches were presented at the same locations (*R* = 0.00, *SE* = 0.10, *CI* = 0.00 – 0.34, *P* = 0.50, *N* = 114; Fig. 1). This suggests that the individual repeatability we observed in food patch visitation was not merely a consequence of individual differences in movement behaviour.

**Figure 1.**
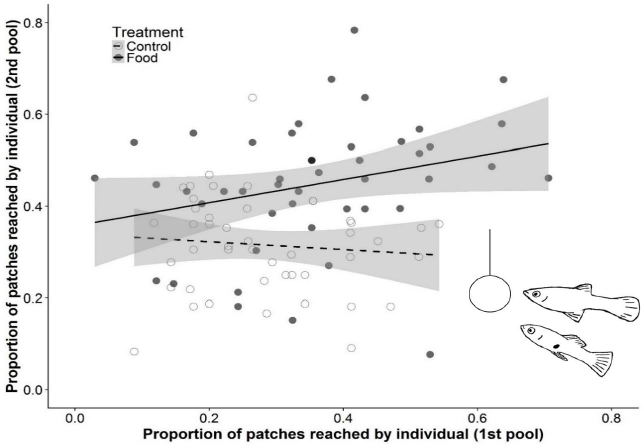
The relationship between the proportion of food and control patches reached by an individual fish in its initial pool (1st pool) and after translocation (2nd pool). Fish showed consistent individual differences in the proportion of food patches reached between pools (solid marker, solid line), but not in control patches reached (open marker, dashed line). Proportions are calculated relative to the total number of food or control patches detected by the batch. Regression lines and 95% *CI* (shaded area) are based on fitted values for proportion of patches reached in the 2nd pool against the 1st pool.

### Individual social tendency, sex and foraging success

More social fish, i.e., fish with a stronger propensity to spend time near conspecifics (before the foraging trials; Fig. 2), reached more patches than less social fish and this effect was stronger for food than control patches (Social time, *10,000 randomization steps*, Food: *coefficient* = 0.20; *P* = 0.001, *N* = 107; Control: *coefficien*t = 0.05; *P* = 0.07*, N* = 107; Fig. 3). Furthermore, males reached more food patches, but not control patches, than females (Sex*Treatment: *χ^2^* = 9.02, *P* = 0.003, *N* = 214; pairwise contrast of females to males during Food treatment: *Z ratio* = -5.31, *P* < 0.001; Control treatment: *Z ratio* = -2.08, *P* = 0.16; Fig. 4). When Sex and Social time were accounted for, individuals were no longer significantly repeatable in the proportion of food patches they reached across environments (Sex: *R* = 0.10, *SE* = 0.13, *CI* = 0.00 – 0.43, *P* = 0.28, *N* = 107; Social time: *R* = 0.26, *SE* = 0.14, *CI* = 0.00 – 0.54, *N* = 114, *P* = 0.07; Sex & Social time: *R* = 0.00, *SE* = 0.10, *CI* = 0.00 – 0.32, *P* = 0.50, *N* = 107), emphasizing the importance of both traits in explaining individual variation in locating food patches.

**Figure 2.**
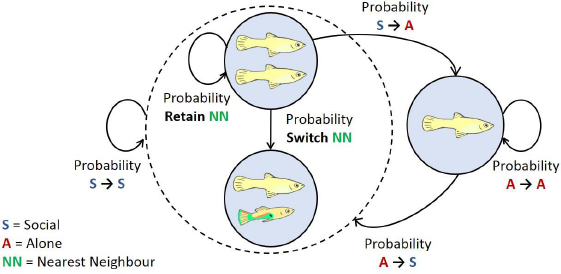
Markov chain model of the guppy fission-fusion social system. When an individual is alone it can, in the next time step, stay in the state ‘Alone’ or go to the state ‘Social’. When in state ‘Social’, an individual can go to the state ‘Alone’ or stay in state ‘Social’. Within a social state an individual can stay with its current nearest neighbour or switch to another. The individual transition probabilities between these states were used to calculate the proportion of time an individual spends near another individual (i.e. in the social state = Social time).

**Figure 3.**
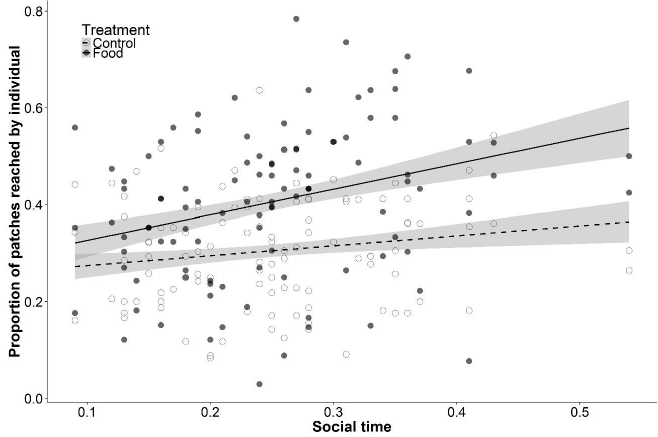
The proportion of food and control patches reached per pool in relation to individual Social time. A higher Social time value indicates a stronger propensity to spend time in proximity of conspecifics (before the foraging trials). In food treatments (solid marker, solid line), but less in control treatments (open marker, dashed line), fish with more Social time reached more patches. Proportions are calculated relative to the total number of food or control patches detected by the batch. Regression lines and 95% *CI* (shaded area) are based on fitted final model values.

**Figure 4.**
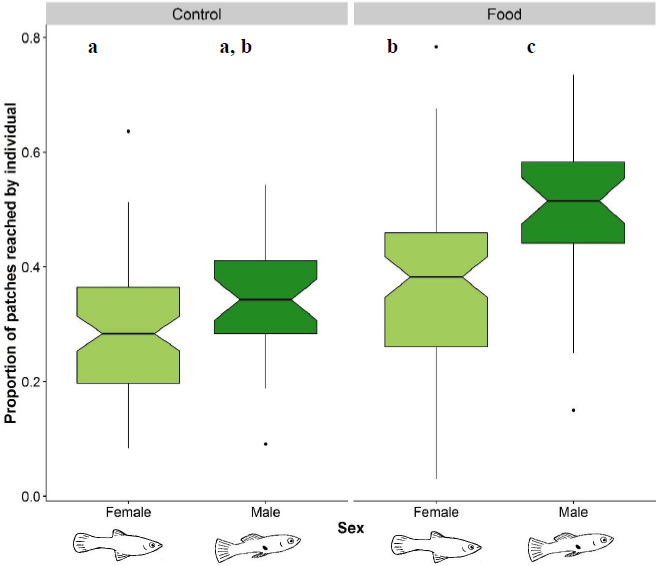
The proportion of food and control patches reached per pool by individual males and females. Males (dark fill) reached more food patches than females (light fill), but not significantly more control patches. Box plots show median and 25th to 75th percentiles with whiskers of 1.5 interquartile distances. Non-overlapping notches suggest a significant difference in medians. Letters ‘a’ to ‘c’ indicate significant differences revealed by post-hoc tests.

That males visited less control than food patches (pairwise contrast of control to food treatment in males: *Z ratio* = −7.28, *P* < 0.001; Fig. 4), suggests that the higher number of food patches reached compared to females was not merely driven by sex-differences in activity. Sex-differences in social behaviour might have played an important role. In line with earlier findings^11^, males and females did not differ in their Social time (within-batch permutation: *10,000 randomization steps*, *difference of means* = 0.02, *P* = 0.46, *N* = 62). Yet, they did differ in the so-called *ϒ*-measure^16^, a measure of the spread of social contact moments across conspecifics (within-batch permutation: *10,000 randomization steps*, *difference of means* = 0.02, *P* = 0.02, *N* = 62), which is not correlated with Social time (within-batch permutation: *10,000 randomization steps*, *coefficient* = -0.06, *P* = 0.39, *N* = 68). That is, males spread their contacts more evenly, while females showed stronger individual preferences for specific social partners. Individuals that spread their social contacts more evenly indeed reached more food and, to a weaker extent, control patches (*ϒ*-measure*, 10,000 randomization steps*, Food: *coefficient* = -0.12, *P* = 0.03, *N* = 107; Control: *coefficient* = -0.04*, P* = 0.048*, N* = 107). Males may thus have reached more food patches than females, partly because they are more flexible in whom they socialize with.

### Number of foraging bites gained

By reaching more food patches, social individuals gained a higher total number of foraging bites than less social individuals (Social time, *10,000 randomization steps, coefficient* = 1.84, *P =* 0.005*, N* = 107), but fish that spread their contacts more evenly did not gain more bites (*ϒ-*measure*, 10,000 randomization steps, coefficient* = 0.69*, P* = 0.83*, N* = 107). Males and females also did not differ significantly in the total number of foraging bites they gained (Sex, *χ^2^* = 3.33, *P* = 0.07, *N* = 107). Females actually tended to have more bites. Females may have compensated their lower proportion of reached patches with a higher bite rate. Indeed, females exhibited a significantly higher bite rate than males when they were present at a patch (Sex: *χ^2^* = 10.13, *P* = 0.001, *N* = 107) and bite rate (similar to the proportion of patch visits) positively correlated with the total number of foraging bites (*χ^2^* = 103.85, *P* < 0.001, *N* = 114). Thus, despite reaching less patches, females did not gain less foraging bites than males. Yet, the strong positive relationship between the proportion of patches reached and the number of bites gained (Supplementary Fig. 2) suggests that if females had reached a similar proportion of food patches to males, they could have gained an even greater number of bites.

### Subpopulation sex ratio and foraging success

Males reached more food patches than females but were equally likely to be the first to arrive at a patch (Supplementary Information; Supplementary Fig. 3). Yet, over time (trial number), males that arrived first at a patch were less likely to be joined by others (Sex*Time: *χ^2^* = 10.99, *P* < 0.001, *N* = 907; Supplementary Fig. 4). Consequently, over time, more members of a batch reached a patch when a female was first compared to when a male was first (Sex*Time: *χ^2^* = 5.59, *P* = 0.02, *N* = 907; Fig. 5). Moreover, individuals were more likely to join at food than at control patches (Proportion of batch joined, Treatment: *χ^2^* = 76.03, *P* < 0.001, *N* = 907). When batch members profit more from a female than a male reaching a food patch, we would expect individuals to experience an increase in foraging success with a relative increase in the number of females. Indeed, individuals (males and females) were overall more likely to reach food patches, but not control patches, the greater the proportion of females in a batch (Sex ratio*Treatment: *χ^2^* = 12.18, *P* < 0.001, *N* = 214; Fig. 6).

**Figure 5.**
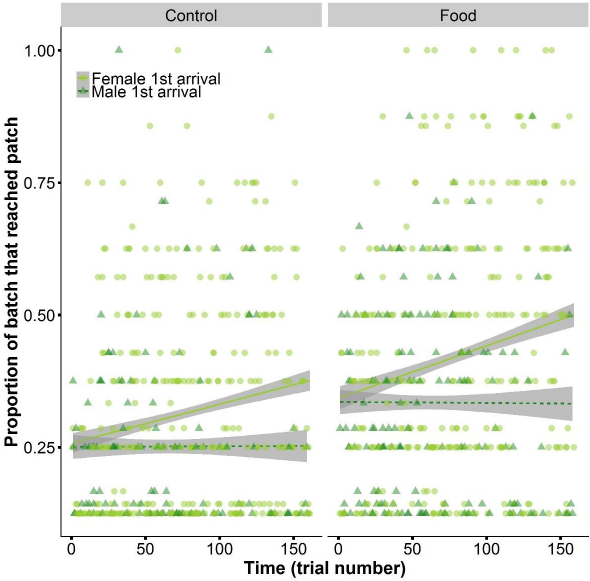
Joining of first arrival by batch members in relation to its sex. Proportion of the adult batch that joins a male (dark triangle) or female (light circle) first arrival at a food or control patch, over time (trial number). Over time, fish were more likely to reach a food or control patch when a female was the first to arrive than when a male was the first. Regression lines and 95% *CI* (shaded area) are based on fitted final model values.

**Figure 6.**
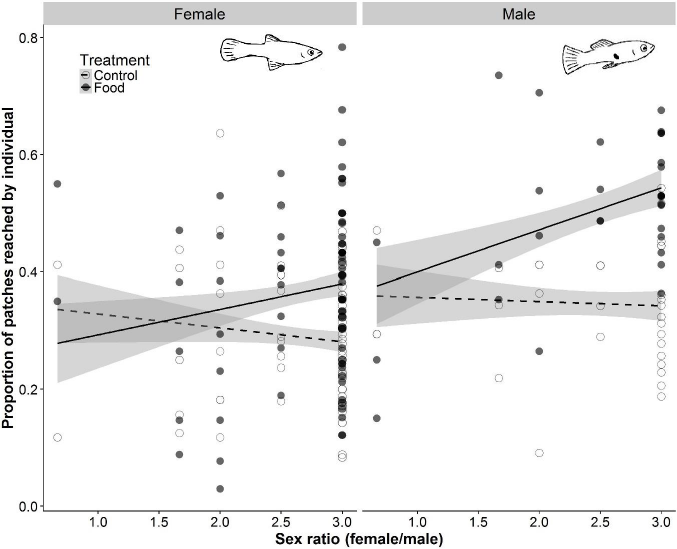
The proportion of detected patches reached by an individual per pool in relation to the sex ratio of its batch. Individuals (males and females) were more likely to arrive at food patches (solid marker, solid line), but not control patches (open marker, dashed line) in batches with stronger female-biased sex ratios. Proportions are calculated relative to the total number of food or control patches detected by the batch. Regression lines and 95% *CI* (shaded area) are based on fitted final model values.

## Discussion

We expected females and more social individuals to be consistently better foragers across environments. Indeed, more social fish reached more food patches than less social fish, yet males were, unexpectedly, more successful in reaching food patches than females. It is intriguing that males reached more food patches than females, since female guppies are considered more food motivated^17,18^. Over time, however, batch members were less likely to join male foragers than female foragers (Supplementary Fig. 4), which might partly explain why, on a subpopulation level, individuals in batches with relatively more females actually reached, on average, more food patches. As males always had one more female to join compared to their female batch members, we expect that this sex-dependent association behaviour was also a key component of the mechanism by which males reached more patches than females.

These findings may have important implications. Social animals typically do not attend to the behaviour of all group members equally^19,20^. As in many animal species^21,22^, male and female guppies both prefer to associate with females over males^23,24^ and as a consequence they could miss foraging opportunities provided by males. Males might ‘attract’ less batch members because they provide weaker foraging cues (e.g. lower bite rate^25^) or because batch members try to avoid male initiated costs (e.g. sexual harassment, male-male aggression)^26–28^. The costs of foregoing or missing foraging opportunities provided by males is probably negligible when natural populations are strongly female-biased^29^. However, based on our findings, when a populations’ sex ratio is 50:50 or male-biased, individuals are expected to experience reduced foraging success. Interestingly, natural guppy sex ratios are known to fluctuate heavily through time (i.e., season) and space^30^, which we predict should have individual foraging consequences. The reduced responsiveness towards foraging males we revealed here could thus turn into maladaptive foraging behaviour and consequently have reproductive consequences for females^31,32^. One possible next step is to test whether individuals in natural populations with primarily equal or male-biased sex ratios have been selected (or have learned) to respond more strongly to foraging males compared to populations with female-biased sex ratios. Alternatively, such a study might reveal that avoiding the costs of being alone with a male outweighs the costs of missed foraging opportunities^31^, irrespective of sex ratio.

Living in groups helps reduce individual predation risk, but is also beneficial when animals forage in unpredictable environments, since it allows individuals to take advantage of the foraging information of others^19,33–35^. Previous studies have indeed identified the social environment as a crucial factor for successful foraging under environmental uncertainty in a wide range of species^36–39^. Through social foraging, animals can obtain information^34,40^ and increase foraging success without the need to individually sample the whole environment^41–43^. Also, the mere presence of others may result in increased foraging success (i.e. social facilitation)^44^. Individual great and blue tits (*Parus major*, *Cyanistes caeruleus*), for example, that joined larger flocks profited from a higher intake of a novel and difficult to acquire food source^45^. And individual barnacle geese (*Branta leucopsis*) which followed information provided by group members benefited from more feeding time^46^.

Our findings that fish reached substantially more food than control patches, and that more food patches were reached by more social fish, strongly indicates that the social environment was indeed an important component in facilitating foraging success. Individuals that have a higher propensity to be social might reach more food patches because they are more attentive to social (foraging) cues^47^ or have an increased close-range exposure to social cues emitted by companions. Being social could also increase the chance that a foraging cue provided by a conspecific at long-range gets noticed, i.e. the many-eyes theory^33^. Not only how much time an individual spends social, also how it distributes its contact moments was a positive predictor of reaching novel food patches. Individuals that spread their social contacts more evenly might likewise spread their social attention more evenly and so more quickly pick up relevant foraging information. This social flexibility partly explained why males reached more food patches than females. Guppies live in a fission-fusion social system, in which they alternate between being alone and associating with one or a few companions. Such a fission-fusion system, which is widespread throughout the animal kingdom, would allow for several of the above mentioned social mechanisms to operate in synergy as so increase individual foraging success.

Both control and food patches in which a female arrived first were, over time, reached by more conspecifics than patches at which a male arrived first. Many of the control patch visitors might have arrived passively by moving together with a companion, i.e. via ‘un-transmitted social effects’^35,48^, but also by responding to ‘false’ cues^49^. More than 56% of the fish that arrived second came within three seconds of the first fish, both at food and control patches. These short latencies suggest that fish were making very fast but potentially inaccurate following decisions (speed-accuracy trade-offs)^50^. For example, acceleration when swimming to a food patch (or mistaken control patch) is likely to be used as a cue. Such a cue may produce many false positives, since it signals what an individual “thinks” it has found, but not what it actually found^51^. In animal populations in which the costs of falsely responding (e.g. increased predation risk, energy loss) are relatively low, conspecific cues might often be wrong^52^, yet right often enough to offset these costs. Alternatively, social individuals might put up with incorrect information just to stay near a companion^49^.

Animals possess several physical and behavioural traits that are consistent over time and/or space^53^. Interestingly, such stable individual traits commonly interact with the social environment^54–56^. Consistent individual traits that interact with potentially important components of foraging, such as the social environment, but also movement patterns and search strategies^1,2^, could generate consistent individual differences in foraging success and so promote selection on these traits. However, to infer selection, it is important to link such traits to consistent foraging success (e.g. finding new resources), independent of the individual’s current environment. In many ecological systems, it is extremely challenging and often unfeasible to completely take individuals out of their environment and place them repeatedly into new ones while remaining under the selective forces of the wild. Here, we took advantage of a key study system in evolutionary ecology, the Trinidadian guppy^57^, and were able to show that individual level traits such as sex and social tendency as well as subpopulation sex ratio can be important drivers of consistent foraging success across different novel environments in the wild.

## Acknowledgements

We are grateful to Sidsel Bouet and Sergio García Martín for assistance with the video analysis and to Félicie Dhellemmes, Herma te Brake and Rieke Seifert for assistance with the data collection. L.S. was funded by an IGB Postdoc Fellowship 2017.

## Author contributions

All authors contributed significantly to the design of the study and the collection of the primary data. L.S. and S.K. analysed the data and L.S. wrote the main manuscript. All authors commented on the manuscript and accepted its last version.

## Additional information

Supplementary material is available for this paper.

## Materials & Correspondence

Correspondence and material requests should be directed to L.S. at

## Methods

### Study area

The study took place in Trinidad in the Upper Turure region (10°41’8”N, 61°10’22”W) from 11 to 30 March 2016 in four natural pools located in a rainforest river system (pool surface area range: 3.1 – 5.6 m^2^, maximum depth range: 15 – 25 cm). The in- and outflow of these pools was slightly altered to reduce fish migration but a continuous water-flow was maintained. All guppies originally occurring in these pools were removed. From nearby pools we caught guppies and individually marked them using an established method of fluorescent elastomer (VIE) colouring^58,59^. We collected two batches of seven and seven batches of eight fish within the natural range of sex and age compositions^29,30^, including 38-75% females, 13-38% males and 0-50% juveniles; comprising a total of 45 females, 19 males and 6 juveniles. Because a few fish escaped after the observations of social behaviour, we finished off with one batch of six, two batches of seven and six batches of eight fish in the foraging trials. After marking, fish were released in the study pool and kept overnight to recover. We caught all fish within one batch from the same downstream pool to assure maximum familiarity of fish within a batch. We performed all research in accordance with the law and animal ethical standards of the country of study, Trinidad and Tobago.

### Social phenotypes

To quantify the social phenotypes, we performed focal follow observations between 09:00 and 15:00. Each fish was followed for 2 min, recording its nearest neighbours every 10 sec (see also^9^). A fish was considered a neighbour if it was within four body lengths of the focal fish^9–11^. After following each fish in a batch for two min, we waited for 10 min to ensure independence of focal sessions^9^, upon which we repeated the procedure. This procedure was repeated 12 times for each batch over two or three days (depending on weather conditions), resulting in a total of 24 min of focal follows per fish in each batch. To quantify an individual’s propensity to be social, we used Markov Chain analysis (see below) to calculate the proportion of time an individual spends near other individuals (i.e. Social time^9,10^). To quantify the degree to which individuals have social preferences, we calculated the *ϒ*-measure as the sum of squares of the normalized association strengths (relative number of contact moments) between one individual and all others^11^. In previous studies with Trinidadian guppies, these social measures have been revealed to be consistent throughout habitat alterations and translocations^10,11^.

### Foraging experiment

To study how sex and social traits influence foraging success we conducted food provisioning experiments. As novel food source we used a small lead ball (8 mm diameter) covered in a mix of gelatine and fish food (TetraMin©). This food item was gently lowered in the pool using monofilament fishing line attached to a rod. Once in the water, the food item was kept approximately 5 cm above the bottom. Upon detection by a guppy, the item was gently lowered to the bottom, with exception of the first trials in the first batches. As control treatment, we used an identical procedure except that the lead balls were not covered with food. These food and control presentations mimic natural events of either food (e.g., insects, fruits) or non-edible items (e.g., leaves, twigs) falling on the water surface and slowly sinking to the bottom, being available for only a limited time^60^. We presented the food and control items in pre-determined feeding locations (zones). In pools 1 to 3 we created ten feedings locations and in pool 4 six feeding locations (because this pool was substantially smaller), assuring roughly equal distance between all feeding locations. We presented control and food items at each location in a randomized order, with the constraint that a location was not used twice in a row. After presenting an item, we waited for a fish to detect it (defined as approaching the item within two body lengths). Upon detection, the item was left in the water for 1 min after which we removed it and the trial ended. If the item was not detected within 3 min the trial also ended. After finishing a trial, we waited for 3 min before starting a new trial. After presenting a food and control treatment at each location, we waited for 30 min upon which we started a new sequence. We performed four such sequences for fish in pools 1 to 3 (over a period of two or three days depending on weather conditions), resulting in 40 food and 40 control trials per batch per pool. In pool 4, which had only six feeding locations, we performed this sequence seven times, resulting in 42 food and 42 control trials per batch per pool. Six of the nine batches of fish were, after their respective foraging trials, caught and relocated to another study pool. The next day, we repeated the foraging experiment in the new pool to study if the observed foraging success was consistent across environments. Two times during the study, an entire batch had emigrated out of the study pool, most likely because of heavy overnight rain, reducing the number of foraging trials in comparison to the other batches (see Supplementary Tables 1-2 for more details on the study time line and batch compositions). In total, we conducted 1,141 trials (incl. one replicate trial).

### Video observations

We recorded all trials with Panasonic and Sony HD Handycams mounted on tripods. From these recordings, we scored the identity, order and feeding behaviour of each newly arrived individual for the 1 min following initial discovery. When a patch remained undetected, we recorded for a maximum of 3 min. Six trials were excluded because the observation time was too short (< 3 min) to reliably quantify a patch as un-detected, leaving 1,135 trials with binary data (yes/no) on patch detection (of which 94 patches/trials were not detected). Due to poor video quality (e.g., water surface glare), some videos were not or only partly useable (e.g. arrival first fish). We could determine the identity of all visiting fish for 963 videos and the feeding behaviour of those fish (e.g. time spent within two body lengths of the (food) item, number of bites taken from the item) for 944 videos. We analysed the videos using the open-source event-logging software BORIS^61^ (v 4.0). For each detected patch, we recorded for 1 min the following variables for each individual fish arriving at the patch: arrival time, duration present (i.e., within two body lengths of the item) and number of bites at the item. Fish identification during the video analysis was cross-validated with the identities reported in the field notes. Two observers analysed all of the videos and showed high inter-observer agreement in individual identification and behaviour (Supplementary Information).

### Statistical analysis

To analyse our foraging experiments, we ran general and generalized mixed models (LMM & GLMM) with R^62^ version 3.4.1 in R Studio version 1.0.153 (© 2009-2017 RStudio, Inc.), using the *lmer* and *glmer* functions in the ‘*lme4’* package^63^. Variables of specific interest (e.g. Sex, Social time) and control variables inherent to the research design were kept in the model at all times, also when they remained non-significant in the final model. These control variables included: Treatment (Control/Food), Relocation (1^st^/2^nd^ pool), Pool identity (Pool 1 to 4), Zone identity (36 patch locations nested in Pool identity), Batch identity (9 batches) and Fish identity (62 adults and six juveniles nested in Batch identity). Because the sex of juvenile guppies could not be reliably determined, models including Sex excluded data for juveniles (3% of the data).

We always started with full models, containing all variables (see Supplementary Table 3 for an overview). To test the significance of fixed effects, we compared models with and without the fixed effect of interest, using Log Likelihood Ratio (LLR) tests. Fixed effects with *P* > 0.1, that were not variables of interest or the above-mentioned control variables, were removed from the model starting with the highest-level interactions. All continuous variables were centred and scaled. We evaluated model fit of linear models via visual inspection of the fitted versus residual plot and the residual frequency distribution. Binomial models (proportions) were tested for over-dispersion. For further information on model validation, see the Supplementary Information and see Supplementary Tables 4 to 11 for the final model statistics. We based conclusions for social phenotypes on permutation models (see ‘analysis of social effects’).

#### Treatment detection

To test if there was an effect of treatment (control or food) on detection probability, we ran a GLMM (binomial) with patches as unit of analysis (*N* = 1,135). Whether a patch was visited by at least one fish (yes/no) was used as the binary dependent variable. The interaction between Treatment and Time (i.e. trial number, continuous), item Drop after first arrival (yes/no) and Relocation were included in the full model as fixed effects. Batch identity and Zone identity (nested in Pool identity) were included as random effects (Supplementary Table 4). See Supplementary Table 3: ‘model 1’ for model details.

#### Consistent individual foraging success across environments

We quantified foraging success as the number of patches reached by an individual relative to the total number of patches detected by its batch. We calculated foraging success for each Treatment*Pool combination that an individual had experienced, thus resulting in four values for individuals in the six translocated batches and two values for individuals in the other three batches. To test whether foraging success was consistent across environments, we calculated the individual repeatability (*R*) of the proportion of patches reached per pool. We derived repeatability values and their 95% confidence intervals using the ‘rptR’ package^64^. Repeatability was calculated separately for food and control treatments. To assess how much variation in foraging success could be attributed to the individual, repeatability values were calculated based on a model that only included Pool identity and Relocation as fixed effects and Individual identity as random effect. To study the individual drivers of repeatability, we additionally assessed models including the individual traits: Sex and Social time. See Supplementary Table 3: ‘model 2a and b’ for model details.

#### Individual social tendency, sex and sex ratio in relation to foraging success

To test the effect of social tendency and sex on foraging success, we ran a GLMM with the proportion of patches reached as dependent variable (*N* = 214). Sex, Social time, Sex ratio and their interaction with Treatment were added as fixed effects. As a proxy for foraging motivation, Bite rate (centred on Sex, since Sex influenced Bite rate) was added as additional control variable. Pool identity and Relocation were again added as fixed effects and Individual identity (nested within Batch identity) as random effect (Supplementary Table 5). Sex ratio was not correlated to the number of foraging adults in a batch (*Spearman Rho* = 0.38, *P* = 0.31, *N* = 9) and replacing Sex ratio with the absolute number of males or females in a batch did not lead to a better model fit (*∆AIC* = + 4.1 & *∆AIC* = + 7.4, respectively). To test for a potential effect of body size (i.e. Body length (mm)), we ran the final model again, replacing Social time with Body length, since Social time was correlated to Body length (Within-batch permutation, overall correlation coefficient = 0.40, *P* < 0.01, *N* = 68). Body length did not significantly affect the proportion of food or control patches reached (Body length*Treatment: *χ^2^* = 3.78, *P* = 0.052, *N* = 214, but *χ^2^* = 4.06, *P* = 0.044 after removal of a potential outlier; Supplementary Table 6). See Supplementary Table 3: ‘models 3a and b’ and for model details.

#### Total number of gained bites and bite rate

To test the effects of (1) individual foraging behaviours and (2) individual traits on the total number of bites, we ran two models (due to collinearity issues). For each individual, we calculated its total number of bites as the sum of all its bites per pool, divided by the total number of patches detected by its batch per pool (only food treatment). This measure thus expresses the average number of bites acquired by an individual over all patches detected by its batch. For each individual, we also calculated its bite rate as the sum of all its bites per pool, divided by the sum of time (seconds) present at a patch per pool (only food treatment). The first model (LMM) included the Proportion of patches reached and Bite rate (not centred on sex) as independent variables (*N* = 114; Supplementary Table 7) and the second model (LMM) included Sex and Social time (*N* = 107; Supplementary Table 8). Both models included Pool identity and Relocation as fixed effects (not as random effects due to a low number of factor levels) and Individual identity (nested in Batch identity) as random effect. To test for sex differences in Bite rate, we ran an additional third model (LMM) with Sex, Pool identity and Relocation as fixed effects and Individual identity (nested in Batch identity) as random effect (*N* = 107; Supplementary Table 9). See Supplementary Table 3: ‘models 4a and b’ and ‘model 5’ for model details.

#### Likelihood of conspecifics joining at a patch

To test if males and females differed in how likely they were to be joined at a patch, we quantified for each patch whether a first arriving individual was joined the following 60 seconds (yes/no) and calculated what proportion of the batch reached the patch. We ran two GLMM’s (binomial) for both dependent variables, including Sex and Social time and their interactions with Time and Treatment as fixed effects. Relocation and item Drop after first arrival were added as additional fixed effects and Zone identity (nested in Pool identity) and Individual identity (nested in Batch identity) as random effects (*N* = 907 both models; Supplementary Tables 10 and 11). See Supplementary Table 3: ‘model 6 and 7’ for model details.

### Analysis of social effects

#### Markov Chain analysis

We used the Markov chain based fission-fusion model by Wilson *et al.* (2014)^9^ to describe the underlying social dynamics of the observed focal fish and determine the social phenotype of each individual. The social behaviour of each fish is described as a sequence of behavioural (social) states, being either in the presence of a specific conspecific (within four body lengths) or alone. We used the collected observational data to estimate the transition probabilities between each state for each individual fish [see the Supplementary material of Wilson *et al.* (2014)^9^ for more details]. The individual proportion of Social time equals *P*_a→s_ / (*P*_s→a_ + *P*_a→s_), where *P*_a→s_ is the probability of ending being alone and *P*_s→a_ is the probability of ending a social contact and (Fig. 2).

#### Preferred relationships

We analysed the presence of preferred relationships between the individuals using a randomisation test where we permuted the identities of the focal individuals’ contact partners within each batch. We computed the variation coefficient of the association strengths (numbers of contact moments) for each batch and used the sum of these values as our test statistic. The social structures indeed showed evidence of significant individual social preferences within the subpopulations (within-batch permutation: *10,000 randomization steps*, *sum of variation coefficients* = 5.7, *P* < 0.001, *N*=70), making the *ϒ*-measure (relative distribution of social contact moments over social companions) a relevant social measure in this study.

#### Randomization tests

Effects of social traits (Social time and *ϒ*-measure) on the proportion of detected patches and the number of bites were tested by randomizing the social metrics between individuals within a batch and calculating the coefficient for the effect of the social trait 10,000 times. The original coefficient in the final model was then compared to the distribution of the coefficients of the permutated final models^65^. We conducted this procedure separately for food and control trials. Because the *ϒ*-measure is sensitive for differences in batch size, we ran the analyses with *ϒ-*measure values corrected for batch size (see Supplementary Information). Details of the final models can be found in the Supplementary Tables 5 and 8. To test if the effect of Sex on foraging success could be explained by males spreading their contacts more evenly (smaller *ϒ-*measure value, see Results), we replaced Sex in the models with *ϒ*-measure.

To analyse the influence of Sex on Social time, we permuted the individual Social times within each batch and used as a test statistic the absolute value of the difference of the mean Social times between males and females. We analysed the influence of Sex on the *ϒ*-measure in the same way. To analyse the relationship between Social time and *ϒ*-measure, we permuted the individual Social times within each batch and used as a test statistic Pearson’s correlation coefficient between Social time and *ϒ*-values. Similarly, we analysed the connection between Social time and Body length. Social time and *ϒ*-measure were computed based on the complete batches (*N* = 70 individuals). For our tests, however, we only used those (adult) individuals that were present in the foraging trials (*N* = 62). Also, as in the above described tests regarding the effect of social traits, we ran these analyses with *ϒ*-measure values corrected for batch size (see Supplementary Information).

### Data availability

The datasets analysed in this study are available from the corresponding author on request.

